# Architecture of the ATP-driven motor for protein import into chloroplasts

**DOI:** 10.1101/2024.09.02.610899

**Authors:** Ning Wang, Jiale Xing, Xiaodong Su, Junting Pan, Hui Chen, Lifang Shi, Long Si, Wenqiang Yang, Mei Li

## Abstract

Thousands of nuclear-encoded proteins are transported into chloroplasts through the TOC-TIC translocon spanning the chloroplast envelope membranes. A motor complex pulls the translocated proteins out of the TOC-TIC complex into the chloroplast stroma by hydrolyzing ATP. The Orf2971-FtsHi complex was suggested to serve as the ATP-hydrolyzing motor in *Chlamydomonas reinhardtii*, but little is known about its architecture and assembly. Here, we report the 3.2-Å resolution structure of the *Chlamydomonas* Orf2971-FtsHi complex. The 20-subunit complex spans the chloroplast inner envelope with two bulky modules protruding into the intermembrane space and stromal matrix. Six subunits form a hetero-hexamer potentially providing the pulling force through ATP hydrolysis. The remaining subunits, including potential enzymes/chaperones, likely facilitate the complex assembly and regulate its proper function. Our results provide the structural foundation for mechanistic understanding of chloroplast protein translocation.

## Introduction

Chloroplasts are organelles responsible for photosynthesis, and they are exclusively found in the cells of plants and eukaryotic algae. Chloroplasts presumably originated from ancient cyanobacteria, and they were formed through an endosymbiotic event approximately 1.2 billion years ago (Margulis, 1975; Stiller, 2007; Zimorski et al., 2014). Over time, most cyanobacterial genes were transferred to the nucleus. As a consequence, proteins required in the chloroplast but encoded by nuclear genes need to be transported back into chloroplasts. This process is highly conserved and well-coordinated, and mediated by a protein translocation machinery, which comprises the TOC and TIC complexes located at the outer and inner envelope membranes (Da Been Kim et al., 2023; Hristou et al., 2020; Kikuchi et al., 2018; Li and Chiu, 2010; Rochaix, 2022). The TOC components contain GTP-binding domains which facilitate the recognition and delivery of proteins across the TOC complex. In comparison, the translocation process of proteins across the TIC complex is driven by energy derived from ATP hydrolysis catalyzed by a motor complex. Most of these nucleus-encoded chloroplast proteins contain at their N-terminal end a transit peptide which is cleaved in most cases upon its emergence from the TIC complex into the chloroplast stroma.

Remarkable advances have been achieved in the field of chloroplast protein translocation (Ling et al., 2019; Zhao et al., 2022). Nevertheless, the precise composition of the TIC complex remains to be defined (Nakai, 2020). While it was previously thought that Tic20, Tic40, and Tic110 constitute TIC components responsible for protein import at the inner envelope membrane (Kovács-Bogdán et al., 2010), recent studies demonstrated that Tic20, Tic56, Tic100 and Tic214 form a TIC complex that is responsible for importing proteins in *Arabidopsis thaliana* (Arabidopsis throughout) and *Chlamydomonas reinhardtii* (Chlamydomonas throughout) (Kikuchi et al., 2013; Ramundo et al., 2020). In agreement with these findings, several studies also showed that Tic40 and Tic110 are absent in the TOC-TIC complex purified through affinity chromatography from transgenic Chlamydomonas strains that contain affinity-tagged Tic20, the consensus component of the two types of TIC complexes (Jin et al., 2022; Kikuchi et al., 2018; Liu et al., 2023; Ramundo et al., 2020). However, the relationship between the two independently characterized TIC complexes remains unclear.

Another debate concerns the identity of the proteins of the TIC-associated ATP-driven import motor involved in chloroplast protein transport (Li et al., 2020; Nakai, 2020). Previous studies showed that the Hsp93 and cpHsp70 chaperones provide energy to facilitate protein transport and form a complex with Tic110 and Tic40, while Hsp90C acts as an additional import motor component (Li et al., 2020). Recently, a 2 MDa inner envelope membrane-bound ATPase motor was identified in Arabidopsis (Kikuchi et al., 2018). The complex was shown to contain the plastid-encoded protein Ycf2 and five nucleus-encoded FtsH(-like) proteins and was thus termed Ycf2-FtsHi (FtsH-inactive). Ycf2 was co-purified with the TIC component Tic214, indicating that the Ycf2-FtsHi complex, but not Hsp chaperones as previously proposed, acts as an import ATPase motor for this type of TIC complex (Kikuchi et al., 2018). Similar results were obtained for the TIC-associated ATPase motor of Chlamydomonas (Ramundo et al., 2020; Xing et al., 2022). It was shown that the plastid-encoded protein Orf2971, the Chlamydomonas ortholog of Ycf2, is co-purified with the TOC-TIC complex, together with several other FtsH-like proteins (Ramundo et al., 2020). Moreover, expression of Orf2971 could be correlated with the expression of the key TIC component Tic214 (Ramundo et al., 2020). A recent study showed that Orf2971 is part of a TIC-associated complex in Chlamydomonas (Jin et al., 2022). Depletion of Orf2971 destabilized the TIC complex, resulting in preprotein accumulation in the cytosol (Xing et al., 2022). These results suggest that the Orf2971-FtsHi complex functions as the protein import motor in Chlamydomonas.

Recently, cryo-EM structures of the Chlamydomonas TOC-TIC complex were solved (Jin et al., 2022; Liu et al., 2023), yielding critical new insights into the mechanistic details of how proteins translocate across the two chloroplast envelope membranes. These structures identified the major subunits of TOC-TIC, and confirmed that subunits Tic20, Tic56, Tic100 and Tic214 are constituents of the TIC complex. While several components of the Orf2971-FtsHi ATPase motor were identified by mass spectrometric analysis of the purified TOC-TIC complex, the actual motor complex was missing in the cryo-EM structure. To obtain a more detailed understanding of the mechanism of protein import, solving the structure of the ATPase motor is pivotal. Here, we report the high-resolution cryo-EM structure of the Chlamydomonas Orf2971-FtsHi motor complex. Our results provide novel insights into the molecular mechanism responsible for motor complex assembly and protein translocation.

## Results

### Overall structure of Orf2971-FtsHi

To determine the composition and structure of the authentic ATPase motor complex in Chlamydomonas, we purified the Orf2971-FtsHi complex from the Chlamydomonas strain containing the Flag-tagged Orf2971 as previously described (Xing et al., 2022), and solved its cryo-EM structure at an overall resolution of 3.2 Å (Fig. 1, Fig. S1, Table S1, Movie S1). The complex has a vertical dimension of approximate 210 Å, and it reaches out to the intermembrane space (IMS), spans the inner envelope membrane (IEM) and protrudes into the chloroplast stromal matrix. Based on our high-quality cryo-EM densities, 20 protein subunits have been identified and assigned in the Orf2971-FtsHi structure (Fig. S2, Table S2). The presence of these proteins was further confirmed by our mass spectrometric analysis of the purified complex. Moreover, several components from the TOC-TIC complex were co-purified with the ATPase motor (Fig. S1A), indicating that Orf2971-FtsHi is closely associated with the TOC-TIC complex.

**Figure 1.**
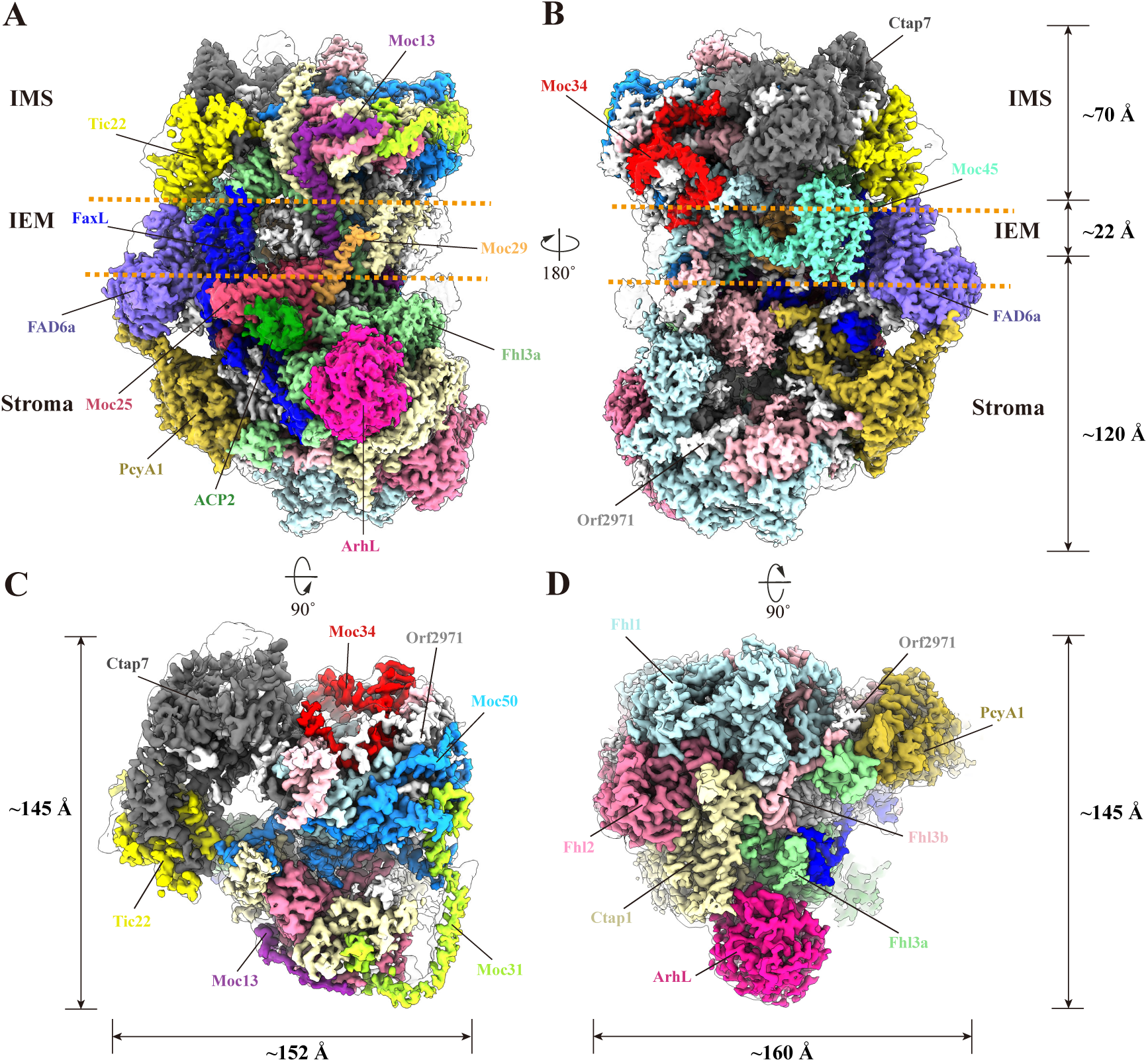
The overall structure of the Orf2971-FtsHi motor complex. **A-B**, Side views of Orf2971-FtsHi motor complex along inner membrane plane (IEM, represented by the orange dotted lines). The protein subunits are painted in different colors. IMS, intermembrane space. **C**, Top view of Orf2971-FtsHi motor complex from the side of the intermembrane space. **D**, Bottom view of the Orf2971 motor complex from the stromal side.

In addition to Orf2971, which is evolutionarily related to the FtsH protein (Kikuchi et al., 2018) and was shown to exhibit ATPase activity (Xing et al., 2022), we identified five other FtsH-like proteins in the ATPase motor structure, including Ctap1, Fhl1 and two copies of Fhl3 (named Fhl3a and Fhl3b). While these proteins were previously suggested to be components of the ATPase motor (Ramundo et al., 2020), the fifth protein is a previously unidentified protein belonging to the FtsH family, based on our Pfam search (Fig. S3A). We thus named this protein Fhl2. An earlier report identified several translocon-related proteins in Chlamydomonas, including Ctap1-7 (Ramundo et al., 2020). Among these proteins, Ctap3-5 were previously shown to be TOC-TIC components, while Ctap1 and Ctap7 were identified here in our motor structure. Notably, our analysis showed that the amino acid sequence of the N-terminal region (1-418) of Fhl2 is identical to that of Ctap6, which contains 446 amino acids (Fig. S3B). On the basis of this finding, we hypothesized that the previously identified Ctap6 (Ramundo et al., 2020) corresponds to the N-terminal fragment of Fhl2 and belongs to the motor complex.

Among the remaining 14 subunits, two proteins named Ctap7 and Tic22 were previously identified as translocon-associated proteins (Kouranov et al., 1998; Kouranov and Schnell, 1997; Ramundo et al., 2020; Rudolf et al., 2013). Both proteins are located in the intermembrane region of the complex (Fig. 1). Sequence and structural analyses of other identified protein components revealed that five subunits are similar to known proteins. Three of these five proteins are located in the stromal region of the complex (Fig. 1A). One protein called phycocyanobilin ferredoxin oxidoreductase (PcyA1) was shown to catalyze the production of phycocyanobilin (PCB), thus appears to function in light sensing. Another protein adopts a structural fold similar to ADP-ribosylglycohydrolase (ARH). We thus termed it ARH-like protein (ArhL). The third protein represents an acyl carrier protein (ACP2) that functions in fatty acid synthesis. Two other proteins are embedded in the inner envelope membrane (Fig. 1A). One subunit named FAD6a was previously annotated as a μ-6 fatty acid desaturase (FAD6)-like protein. Another protein exhibits an overall folding resembling fatty acid exporter (FAX), and was thus named FAX-like protein (FaxL). The additional seven subunits adopt a fold with no detectable similarity to known proteins. We thus named these uncharacterized proteins according to their molecular mass using the prefix Moc (motor of chloroplast) (Table S2).

Previous studies suggested that the plastid NAD-dependent malate dehydrogenase (pdMDH) in Arabidopsis stably associates with the Ycf2-FtsHi motor, and plays an essential role in preprotein translocation (Kikuchi et al., 2018; Schreier et al., 2018). Although it was identified in our Orf2971-FtsHi complex using mass spectrometry (Fig. S1A), we failed to identify any densities that corresponded to the pdMDH, suggesting that Chlamydomonas pdMDH may only loosely bind to the Orf2971-FtsHi motor.

### Core subunits of Orf2971-FtsHi and the hexameric assembly of their stromal domains

Our structure analysis showed that Orf2971 and five FtsH-like proteins are the largest proteins in the ATPase motor. These six proteins account for more than half of the size/molecular mass of the motor complex (Fig. 2A, Table S2). Furthermore, these subunits are the only six subunits that contain an ATPase domain (Fig. S4). Together, these six subunits constitute the core of the motor and form a hexameric ATPase module. When analyzing the structure of each individual core subunit, we found that they share a number of similarities, both in terms of structure as well as topology. They are all membrane-bound proteins whose N- and C-terminal domain (NTD and CTD) are located in the intermembrane and stromal regions, respectively (Fig. 2A,B, Fig. S5). While their NTDs are folded in significantly different ways, they all contain the classical FtsH periplasmic domain (PD^FtsH^) except Orf2971 (at least not in the modeled part in our structure) (Fig. S6A). In contrast, their CTDs are largely similar (Fig. S6B), all containing a typical FtsH cytoplasmic domain (CD^FtsH^), which is usually composed of an AAA^+^ (ATPases associated with diverse cellular activities) domain followed by a protease domain. The AAA^+^ domain in FtsH proteins serves as an engine-like module that converts ATP hydrolysis into mechanical force. This force is then used to transfer the unfolded polypeptide to the adjacent protease domain for further proteolytic cleavage (Suno et al., 2006). Previous studies analyzing the structures of AAA^+^ proteins showed that the ATPase domain consists of a large and a small subdomain (Puchades et al., 2020; Suno et al., 2006). The large subdomain harbors structural motifs essential for ATP binding and hydrolysis, and is linked to the protease domain through the small subdomain. In our structure, we modeled the CTDs of all six core subunits, except for the large subdomain of Ctap1 and the small subdomain of Fhl3b (Fig. S6B). These subdomains exhibit weak densities, presumably due to their conformational variation, and were low-pass filtered at approximately 10 Å to show the general appearance. In addition, the zinc-binding motif HEXXH critical for protease activity (Mishra and Funk, 2021) is lacking in the protease domain of all core subunits (Fig. S4). Thus, while the hexameric core is able to hydrolyze ATP, it is unable to cleave polypeptides. In summary, our structural analysis of the Chlamydomonas Orf2971-FtsHi complex reveals features similar to the Arabidopsis Ycf2-FtsHi motor that was suggested to have originated from the ancestral membrane-bound hexameric FtsH protease and does not require the protease activity for the protein import function (Kikuchi et al., 2018).

**Figure 2.**
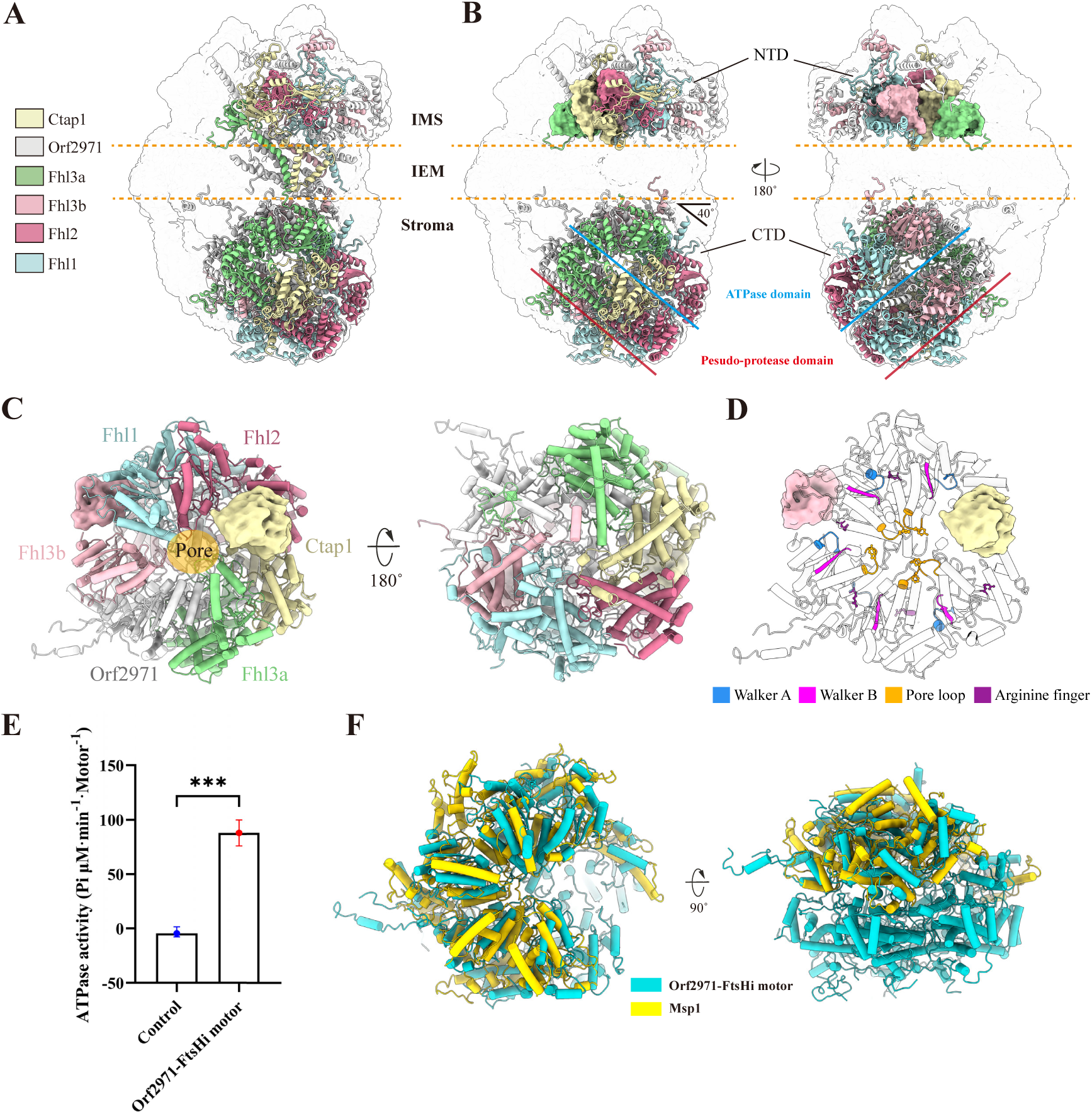
Core subunits of Orf2971-FtsHi motor. **A**, Cartoon representation of the six core subunits wrapped by a transparent gaussian lowpass filtered density. **B**, The NTDs and CTDs (including the ATPase and pseudo-protease domains) of the six core subunits. The five PD^FtsH^ domains are indicated in surface style. The PDs^FtsH^ of Ctap1 and Fhl3a are close to the intermembrane surface of IEM, whereas PDs^FtsH^ of Fhl1, Fhl2 and Fhl3b are away from the IEM. **C**, Top (left) and bottom (right) views of the hexameric assembly of CTDs of six core subunits, showing the ATPase (left) and pseudo-protease (right) domains. The central pore region is indicated. The large subdomain of Ctap1 and small subdomain of Fhl3b were not built due to their weak densities and are shown in low resolution density map generated by gaussian lowpass filter. **D,** Cartoon model of the ATPase hexamer of the motor complex. The Walker A, Walker B, pore loop with the conserved aromatic residue, and Arginine finger are labeled. **E**, The ATPase activity of the Orf2971-FtsHi motor complex is shown in histogram style. All experiments were repeated three times. P value is equal to 0.0003. The enzyme activity unit is Pi µM**·**min^-1^**·**Motor^-1^. **F,** Structural alignment between CTDs of Orf2971-FtsHi motor and Msp1 protein in closed state (PDB code 6PDW).

Our structure shows that, similar to other FtsH motors, the CTDs of the six core subunits form a relatively ordered hexameric unit (Fig. 2C). The six pseudo-protease domains form a disk-shaped assembly located at the bottom of the complex, whereas the ATPase domains sit on top of the pseudo-protease hexamer, and are positioned close to the IEM plane (Fig. 2B, C). We observed a central pore with a minimal diameter of ∼ 8 Å formed by the ATPase domains of the core subunits. The aromatic residue in the pore loop conserved in other ATPases is located at the central pore region (Fig. 2D), thus facilitating the protein substrate threading into the central pore. Other conserved motifs, including the Walker A, Walker B and Arg-finger, are located at similar positions as in other FtsH ATPase motors. These structural features of the Orf2971-FtsHi complex imply that it is active in ATP hydrolysis, which was confirmed by our ATPase enzymatic assay (Fig. 2E). The hexamer is not arranged parallel to the IEM plane; instead it tilts with a crossing angle of approximately 40 degrees (Fig. 2B). A previous report suggested that the AAA^+^ domains tilted relative to the membrane plane create a gap for the protein substrate to enter the central pore of the hexamer (Carvalho et al., 2021).

The ATPase domain of Ctap1 in the motor complex appears mobile, characterized by its less ordered density. The feature of one mobile ATPase protomer was also observed in other AAA^+^ proteins. For example, Msp1 is a transmembrane AAA^+^ ATPase extracting mislocalized tail-anchored proteins from the membrane (Chen et al., 2014; Okreglak and Walter, 2014). The previously reported structure of Msp1 bound with peptide substrate revealed that in its “closed” conformation, one ATPase protomer is mobile and shows dispersed density (Wang et al., 2020). The presence of one mobile ATPase protomer in the FtsH(-like) hexamer is presumably part of the “hand-over-hand” mechanism that is adopted by AAA^+^ proteins to drive protein substrate translocation. Interestingly, the five ATPase domains found in our structure can be aligned with the “closed” Msp1 structure (PDB code 6pdw) (Fig. 2F). Hence, Orf2971-FtsHi may utilize a similar mechanism for pulling the protein across the inner envelope membrane. Analysis of our Orf2971-FtsHi structure shows that in addition to the conserved PD^FtsH^ and CD^FtsH^, all six core proteins contain extra structural elements (Fig. S6A, B). These elements are involved in interactions with other proteins, and are therefore likely to facilitate the assembly and stability of the complex. Moreover, Orf2971 contains multiple additional helices and loops, which are interspersed throughout the complex (Fig. S6C). As a result, Orf2971 forms extensive interactions with 17 of the other 19 subunits present in the complex (Fig. S6D), strongly suggesting that in addition to ATP hydrolysis, Orf2971 functions as a scaffold and plays an essential role in assembling the complex and maintaining its stability. Consistently, previous results demonstrated that Orf2971 is indispensable for cell survival (Xing et al., 2022).

### The IMS module of the Orf2971-FtsHi complex

The Orf2971-FtsHi complex contains a large IMS module comprised of the NTD of six core subunits as well as (the IMS-region of) six auxiliary proteins. The six core subunits and Moc50 (Fig. S7A) together form a compact structure, with the remaining five auxiliary proteins located at the periphery (Fig. 3A). Moc50 contains DnaJ-like zinc finger motifs (Fig. S7B,C), suggesting it may retain functions similar to other DnaJ(-like) proteins, acting as a chaperone or assisting complex assembly. Consistent with this suggestion, our structure revealed that Moc50 is tightly intertwined with the core subunits, suggesting that Moc50 serves as an additional scaffold protein that is critical for facilitating the assembly and stability of the motor complex.

**Figure 3.**
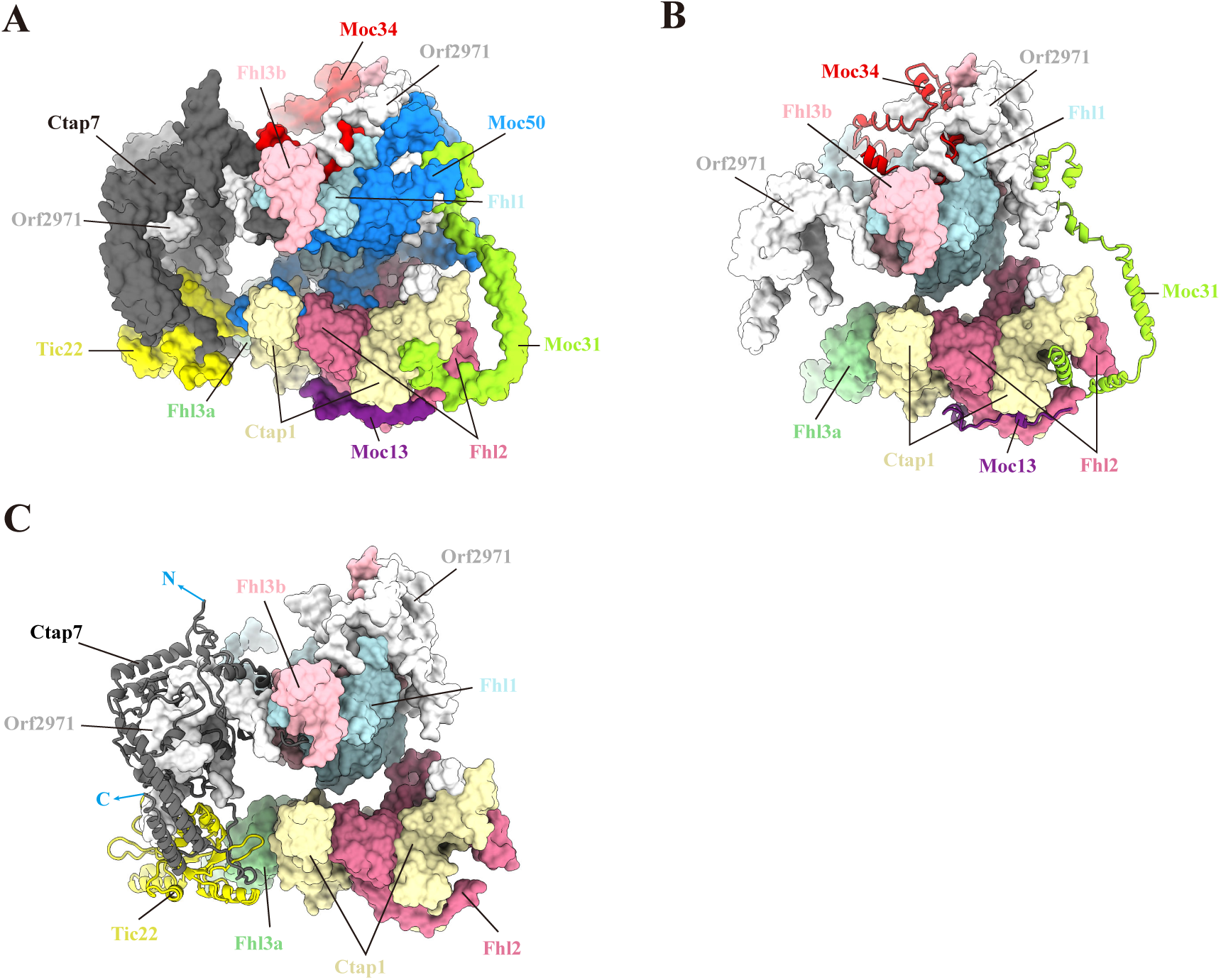
IMS module of Orf2971-FtsHi motor. **A,** Top view of IMS module. **B,** The arrangement of Moc31, Moc34 and Moc13 (shown in cartoon mode) relative to the core subunits (shown in surface mode). **C,** The arrangement of Tic22 and Ctap7 (shown in cartoon mode) relative to the core subunits (shown in surface mode). The N- and C-terminus of Ctap7 are indicated by blue arrows.

Moc31 and Moc34 are two IMS-localized proteins and primarily composed of α-helices. Moc13 possesses one transmembrane helix (TMH) and its N- and C-terminal fragments extend into the intermembrane and stroma regions (Fig. S8). The three proteins together embrace the core subunits from one side, whereas Tic22 and Ctap7 are closely associated and located at the other side of the IMS module (Fig. 3). Tic22 exhibits a partly symmetrical structure of mixed α/β topology, resembling the folding of cyanobacterial Tic22 (AnaTic22, PDB 4EV1) (Fig. S9A,B). Ctap7 contains a central region that adopts a chalcone-isomerase fold commonly found in fatty acid binding proteins (FAPs) such as Arabidopsis FAP1 (AtFAP1) (PDB 4DOO) (Fig. S9C,D) (Ngaki et al., 2012). In addition, we observed extra density which likely belongs to a co-purified endogenous lipid molecule located in a hydrophobic tunnel within Ctap7 (Fig. S9D). The C-terminal extensions of Ctap7 bridges it to Tic22, and both N- and C-terminal regions interact with core subunits (Fig. 3C). These unique interactions allow Tic22 and Ctap7 to form an arc-shaped wall that is tightly connected to core subunits (Fig. 3A,C).

### The IEM module of the Orf2971-FtsHi complex

Remarkably, the Orf2971-FtsHi complex contains at least 30 TMHs spanning the inner envelope membrane and belonging to the core subunits and six auxiliary proteins (Fig. 4A). Among these auxiliary proteins, Moc29, FAD6a, FaxL and Moc45 constitute intrinsic membrane subunits, and the remaining two (Moc50 and Moc13) only contain one single TMH. Moreover, one membrane-attached protein Moc25 closely contacts the IEM from the stromal side although it lacks a TMH (Fig. 4A).

**Figure 4.**
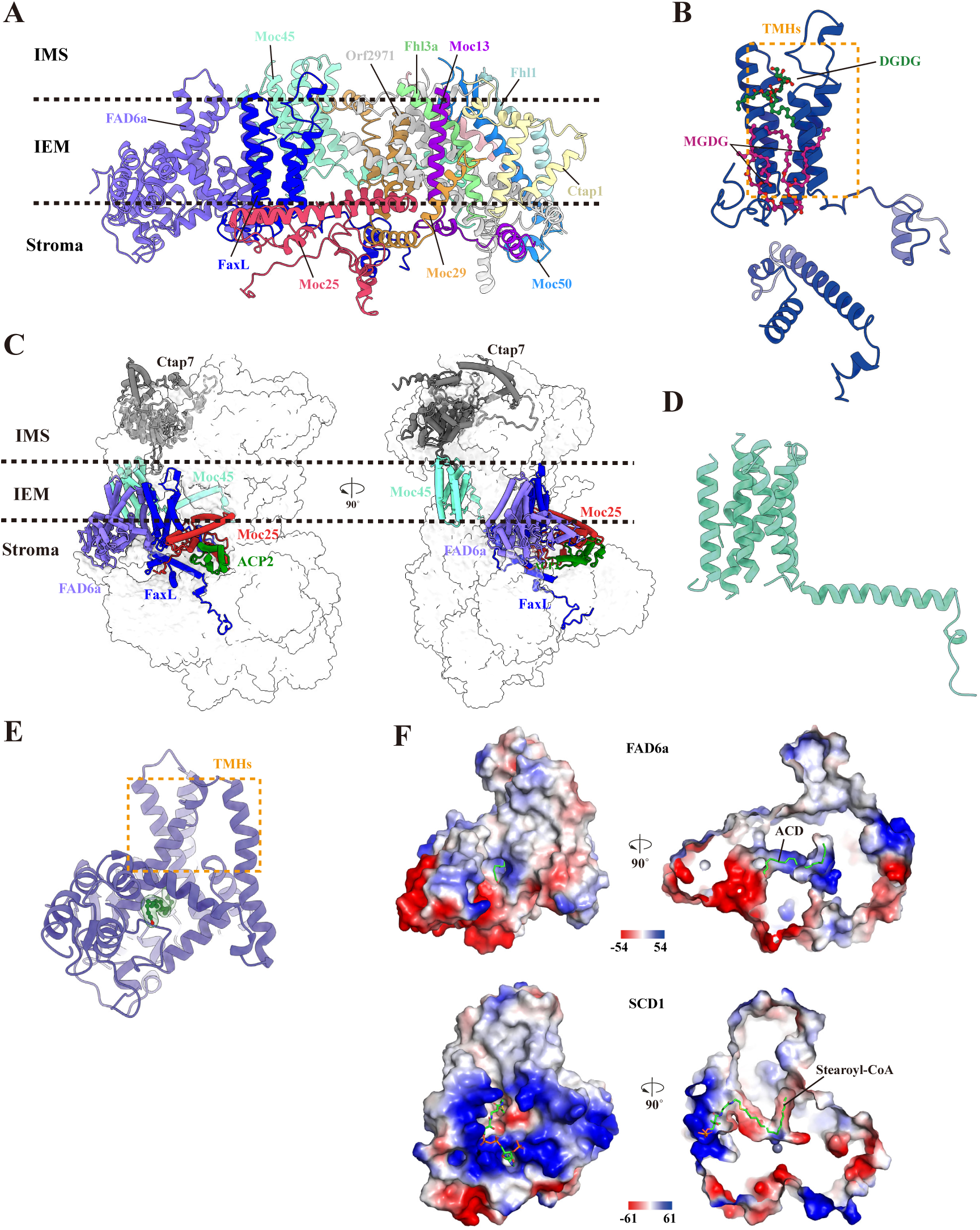
The IEM module and peripherally located proteins. **A,** Transmembrane and membrane-attached proteins are shown in cartoon mode. Boundaries of inner membrane are indicated with black dashed lines. **B,** Overall structure of FaxL. The transmembrane helix region is boxed by an orange dotted rectangle. Lipid molecules are shown in stick mode. **C,** Ctap7, Moc45, FAD6a, FaxL, Moc25 and ACP2 are clustered together and shown in cartoon mode. The black dotted lines represent the inner envelope membrane. **D,** Overall structure of Moc45. **E,** Overall structure of FAD6a. The transmembrane helix region is boxed by an orange dotted rectangle. One arachidonic acid (ACD) molecule was modeled as a putative ligand. The ACD molecule and the cryo-EM density are shown in stick-ball and transparent surface modes. **F**, Electrostatic potential surface and the lipid binding pocket of FAD6a and SCD1. The ligand ACD in FAD6a and stearoyl-CoA in SCD1 are shown as sticks and bound in the corresponding pocket.

Three integral membrane proteins (Moc45, FaxL and FAD6a) are separated from the core, and Moc45 is located more distally from the other two subunits (Fig. 4B-E). We only partially built its structure due to the weak density. Moc45 contains at least seven TMHs and a long C-terminal helix (Fig. 4D, Fig. S10A). FaxL and FAD6a are close to each other. FaxL possesses four TMHs and a stromal N-terminal domain (Fig.4B, Fig. S10B), and is structurally related to the TEMT14 family (IPR005349). It contains a poly-glycine (polyG) motif in its N-terminal domain preceding the four TMHs, this sequence arrangement is similar to Arabidopsis FAX3 (Bugaeva et al., 2023; Li et al., 2015) (Fig. S10C), which was also suggested to be localized in the chloroplast inner membrane, and has a specific N-terminal domain that presumably functions in protein-protein interaction and FA export (Bugaeva et al., 2023; Li et al., 2019; Peter et al., 2022). Consistent with this, we observed several lipid molecules that bind in close vicinity of FaxL (Fig. 4B, Fig. S10D).

FAD6a was annotated to be a homolog of CrFAD6 (Li-Beisson et al., 2015) that belongs to the family of di-iron-containing integral membrane FAD, and adopts a typical FAD fold (Los and Murata, 1998; Shanklin and Cahoon, 1998; Wang et al., 2015; Zhu et al., 2015) (Fig. 4E, Fig. S11A). While we failed to observe the density corresponding to iron ions in FAD6a, three histidine boxes which provide ligands for the di-iron center in membrane FADs were found to be partially conserved in FAD6a (Fig. S11B). Furthermore, we observed a tube-shaped density that fits well with a fatty acid molecule inside FAD6a (Fig. S11E). This FA molecule partially overlaps with the stearoyl-CoA molecule in human stearoyl-CoA desaturase (SCD1, PDB: 4ZYO), which is structurally similar to FAD6a (Fig. 4F).

The remaining three auxiliary proteins (Moc50, Moc13 and Moc29) are clustered together with core subunits (Fig. 4A,5A). Moc29 contains four TMHs and two horizontal helices that are laterally arranged on both sides of the inner membrane (Fig. 5B, Fig. S12A). It folds into a U-shaped structure that secures the transmembrane domain of the core subunits, Moc50 and Moc13 (Fig. 5A,B). Close to Moc29, Moc25 forms a triangular structure with several amphiphilic helices and encloses a number of lipid molecules (Fig. 5C, Fig. S12B-D). One of its helices is parallel to the lateral helix of Moc29 and appears to insert into the inner membrane (Fig. 5D). Taken together, our structural analysis suggests that these yet uncharacterized proteins are critical for the organization of the Orf2971-FtsHi motor complex during assembly.

**Figure 5.**
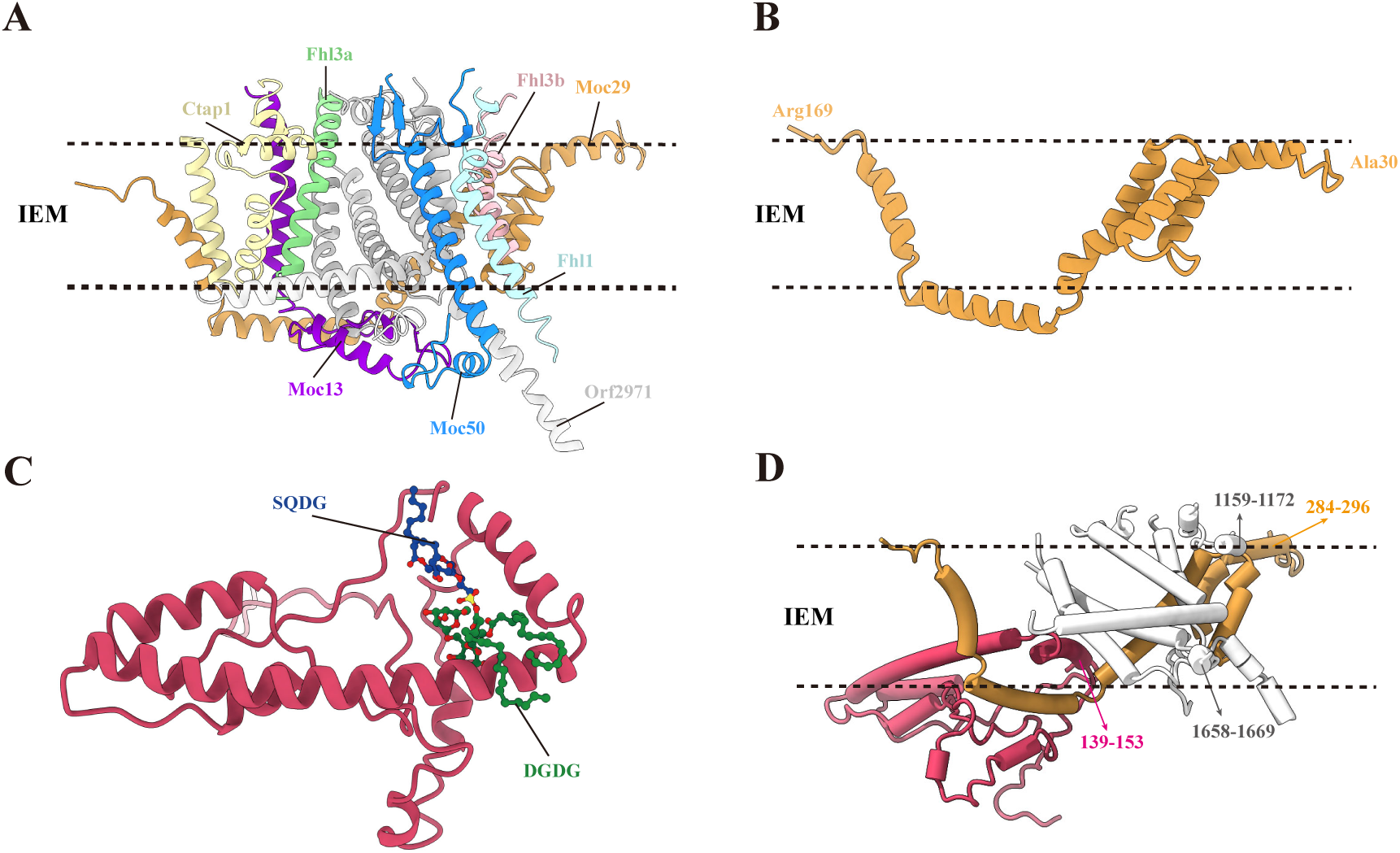
Core subunits and closely related auxiliary subunits in the IEM module. **A,** Transmembrane helices from the core subunits as well as adjacent Moc50 and Moc13 are enclosed by Moc29. **B**, Cartoon model of Moc29 protein shows a U-shaped structure. **C,** Cartoon model of Moc25 protein shows a triangular-shaped structure. Lipid molecules are shown in sticks. **D,** Amphipathic helices from Orf2971, Moc25 and Moc29 confining the vertical dimension of the inner envelope membrane are indicated by arrows. The amino acid numbers that constitute these amphipathic helices are labeled.

Interestingly, we found that Orf2971 also possesses amphipathic helices that may laterally attach to the IEM from both the intermembrane and stroma sides. Together with Moc25 and Moc29 (Fig. 5D), their amphipathic helices confine the vertical dimension of the inner envelope membrane, with a distance of approximately 22 Å. This length is comparable to the thickness of the lipid bilayer membrane. Interestingly, similar feature was also observed in the TIC complex. As shown in previous reports (Jin et al., 2022; Liu et al., 2023), amphipathic helices belonging to Tic20 and Simp3/YlmG of TIC complex are positioned in parallel on the two surfaces of the IEM plane. The presence of amphipathic helices confining the membrane plane in both complexes may help to adjust the membrane thickness by slightly changing the tilting angle of other TMHs, and thus facilitate protein import.

### Three stroma-localized auxiliary proteins

Our structure showed that three auxiliary subunits, namely ACP2, ArhL and PcyA1, are located in the stroma (Fig. 6A). ACP2 adopts a consensus ACP fold containing three parallel α-helices (Chan and Vogel, 2010; Farmer et al., 2019; Li-Beisson et al., 2015), and possesses a DSL motif conserved in other ACPs (Fig. 6B, Fig. S13A-C). In addition, there is an extra density around the hydroxyl group of the conserved serine (pSer72) in ACP2 (Fig. 6B). Comparison of ACP2 in our structure with the previously reported PfACP structure (ACP from *Plasmodium falciparum*, PDB: 3GZL) indicates that the extra density corresponds to the phosphopantetheine (PPT) arm that carries an acyl chain and is covalently linked to the conserved serine residue in ACPs (Agarwal et al., 2012) (Fig. S13B). Although located in the stroma, ACP2 in the complex is embraced by Moc25, which links ACP2 to the inner envelope membrane (Fig. 6A).

**Figure 6.**
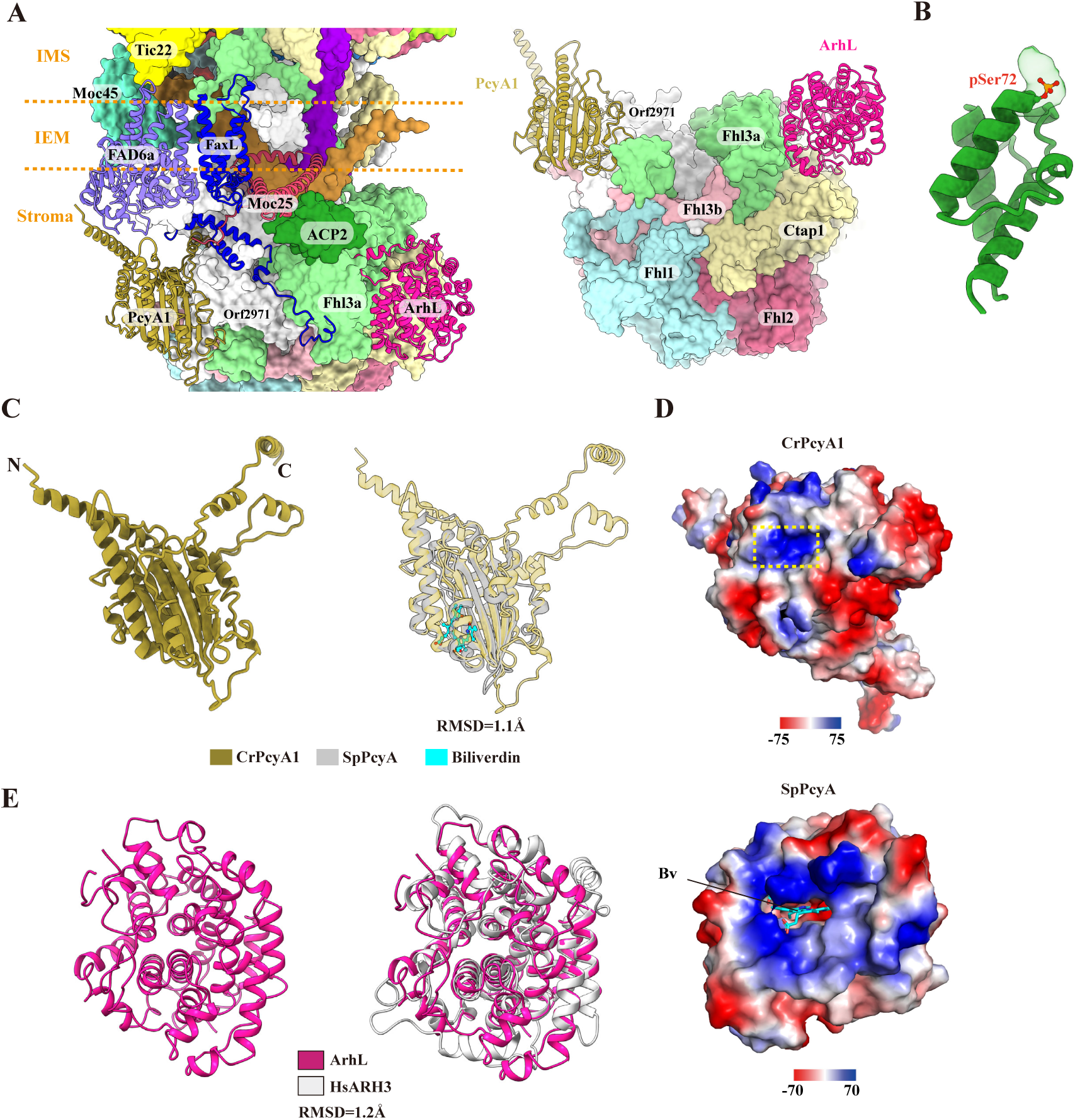
Structure, location and interactions of auxiliary proteins in the stroma. **A,** Binding sites of ACP2, ArhL and PcyA1 within the motor complex, viewed from the membrane plane (left) and from the bottom of the complex (right). PcyA1, ArhL, FAD6a, Moc25 and FaxL are displayed in cartoon mode. Other subunits located in proximity are shown in surface mode. **B,** Overall structure of ACP2. **C,** Overall structure of CrPcyA1 (left) and the structural comparison between CrPcyA1 and cyanobacterial PcyA (SpPcyA) (right). The N- and C-terminus of CrPcyA1 are indicated in the left panel. The biliverdin (Bv) molecule bound to SpPcyA is shown in stick and in cyan color (right). **D,** Electrostatic potential surface of CrPcyA1 and SpPcyA. The Bv binding pocket, which is sealed and exhibits positively-charged surface in CrPcyA1, is highlighted by yellow dashed rectangle. **E,** Overall structure of ArhL (left) and structural comparison between ArhL and human ARH3 (HsARH3) (right).

ArhL and PcyA1 are attached at the periphery of the hexameric CTDs of core subunits (Fig. 6A). ArhL binds to the Ctap1-Fhl3a interface, whereas PcyA1 associates with

Orf2971 and Fhl3a. Their locations suggest that the two proteins are critical for the assembly and stabilization of the core in the stroma. ArhL is an α-helical protein that shows high structural similarity with ADP-ribosyl hydrolase (ARH) 3 (PDB 6D36) and ARH1 (Fig. 6E, Fig. S14D). ARHs hydrolyze and remove the ADP-ribosyl modification, one of the most commonly used post-translational modifications (PTMs) of proteins. However, the key residues for hydrolase activity and magnesium binding in ARH1 and ARH3 are only partially conserved in ArhL (Fig. S14E). Therefore, it is unclear whether ArhL functions as an enzyme similar to ARHs, or whether it is primarily playing a structural role.

PcyA1 exhibits a three-layer α/μ/α sandwich fold (Fig. 6C, Fig. S14A). Structural comparison of CrPcyA1 with its cyanobacterial counterpart SpPcyA (PDB 2D1E) shows that CrPcyA1 possesses N- and C-terminal extensions (Fig. 6C), which contact the membrane-intrinsic subunits FAD6a and FaxL, thus attaching PcyA1 to the inner envelope at the stromal side (Fig. 6A). This finding is in agreement with a previous report showing that CrPcyA1 is partially associated with the membrane of the chloroplast envelope (Duanmu et al., 2013). PcyA1, which was previously identified as a ferredoxin-dependent bilin reductase (FDBR), catalyzes the conversion of PCB from billiverdin (Bv). Our sequence and structural analysis reveals that CrPcyA1 contains conserved residues involved in Bv binding and catalysis (Fig. S14B-C). The Bv pocket of CrPcyA1 has a positively-charged surface and faces the stroma (Fig. 6D), which may allow the binding of negatively-charged ferredoxin. These structural features suggest that CrPcyA1 in the Orf2971-FtsHi complex retains its catalytic activity in addition to its structural role in the complex.

### Translocon-motor model and potential translocation pathways

The precise mechanism of how the TOC-TIC and motor complex cooperate to import preproteins remains to be elucidated. It is likely that direct interactions between the TOC-TIC translocon and the Orf2971-FtsHi motor occur during the translocation process. Previously, Tic22 in the Arabidopsis Ycf2-FtsHi complex was shown to interact with the POTRA domains of AtToc75 (Paila et al., 2016). In line with this previous report, our molecular dynamic simulation showed that Tic22 in the Chlamydomonas Orf2971-FtsHi complex strongly interacts with the POTRA domains of CrToc75 (Fig. S15, Movie S2). We further purified recombinant proteins of CrTic22 and POTRA domains of CrToc75, and measured their binding affinity using surface plasmon resonance. Our data showed that the two proteins can stably bind to each other, with a KD value of 5.22 μM (Fig. 7A). CrTic22 and CrToc75 are both located at the periphery of their corresponding complex (Orf2971-FtsHi motor and TOC-TIC) (Fig. S15A). These two proteins may therefore form the direct contact site between the Orf2971-FtsHi motor and the TOC-TIC translocon (Fig. 7).

**Figure 7.**
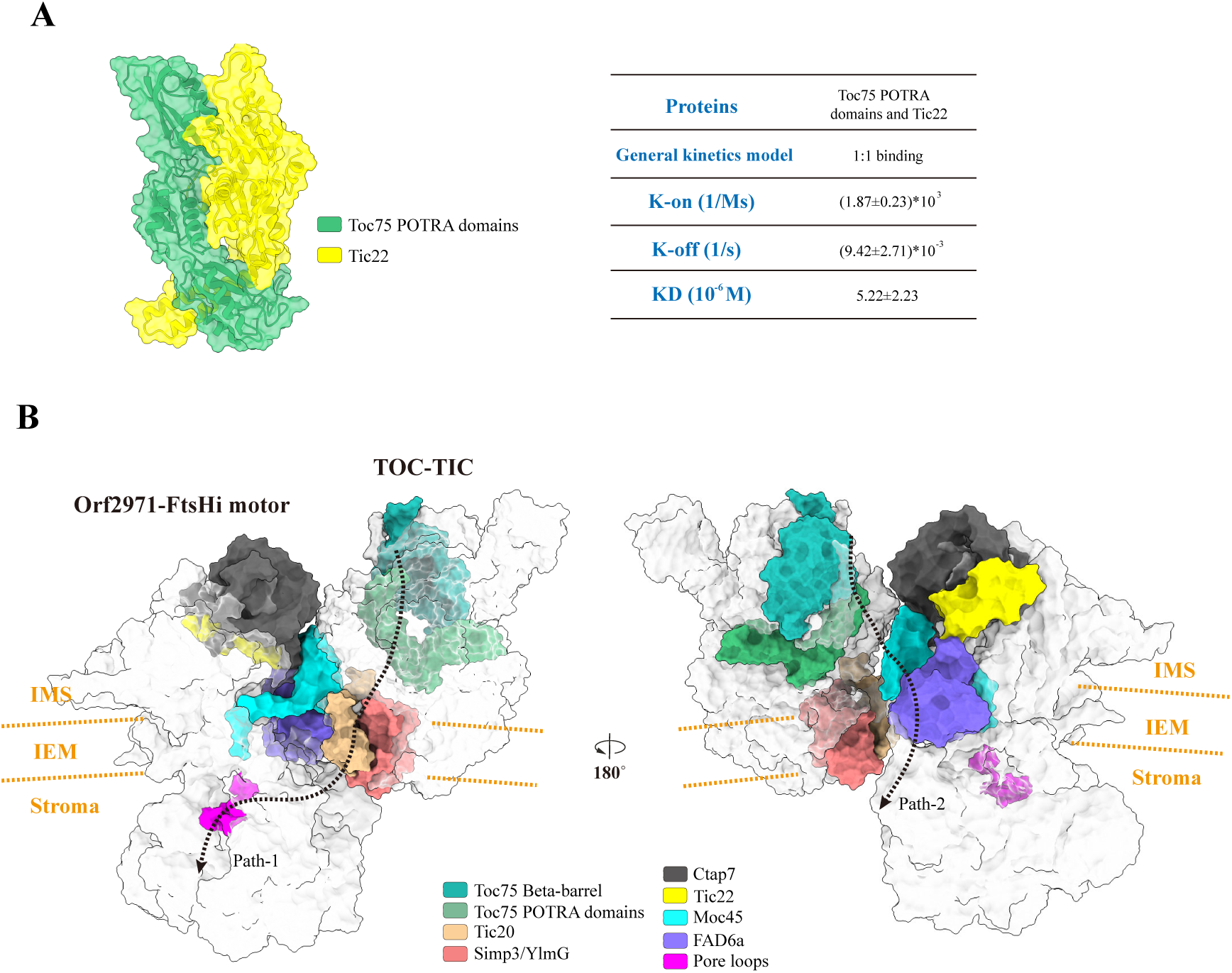
Proposed translocon-motor model. **A,** MD simulation model of POTRA domains of Toc75 bound to Tic22 (left) and their binding affinity measured by SPR experiment (right), shown as the equilibrium dissociation constant (KD). Data are mean ± s.d., calculated from three independent measurements. **B,** Putative interacting model of TOC-TIC translocon and Orf2971-FtsHi motor shown in surface mode. The protein subunits presumably located at the interface between TOC-TIC and Orf2971-FtsHi are painted in different colors. The orange dotted lines represent the inner envelope membrane (IEM). Potential paths for protein translocation are indicated by black lines.

Based on these results, we proposed a hypothetical translocon-motor model, in which Tic22 and Toc75 face each other with short distance (Fig. 7, Movie S1). Tic20, a TIC component which was suggested to constitute a potential path for preprotein translocation, faces towards the motor complex, thus the preprotein imported from the TIC complex can be easily captured by the motor proteins and led to the ATPase central pore (Path-1). In addition, within the motor complex, we observed a wide channel with a lateral opening in the inner envelope, shaped by FAD6a and Moc45 (Fig. 4C), and towards the TIC complex in our proposed model (Fig. 7B). The wide membrane-spanning channel has a diameter of approximately 20 Å, thus could allow a small folded protein to pass. This wide channel, together with the lateral gate formed by Toc75 and Toc90 in TOC-TIC, which was previously proposed to be able to open transiently (Liu et al., 2023), may together form another channel (Path-2). This path may explain the previously reported result that dihydrofolate reductase in Arabidopsis can be imported across chloroplast membrane as a folded protein (Ganesan et al., 2018). Our structural observation of these potential pathways suggests that in addition to hydrolyzing ATP and providing the pulling force for the TOC-TIC complex, the import motor itself may directly participate in the protein translocation process.

## Discussion

Here, we solved the structure of the Orf2971-FtsHi motor at near-atomic resolution, revealing its unique architecture and protein composition. Our structure shows that the CTDs of six core subunits are positioned in the stroma and constitute the ATPase hexamer. These six proteins are involved in hydrolyzing ATP and pulling the protein substrate into its central pore. To provide the pulling force, one would expect that the motor complex is proximal to the TOC-TIC translocon. In agreement with this notion, our binding affinity and MD simulation analysis indicate that Tic22 directly interacts with Toc75, allowing the required interplay between these two complexes during protein translocation. Furthermore, both Tic22 and the POTRA2-3 domains of Toc75 were recently shown to possess chaperone(-like) activity (Glaser et al., 2012; O’Neil et al., 2017), indicating that they cooperate in preventing misfolding of imported proteins. Furthermore, our structure implies that in addition to facilitate the complex assembly, several proteins may also possess specific functions during protein translocation. Earlier reports demonstrated that most of the photosynthetic pigment binding proteins are imported by the TOC-TIC complex (Bauer et al., 2000; Jarvis et al., 1998; Jouhet and Gray, 2009). Pigment molecules are required for proper folding and functionality of these photosynthetic proteins (PAULSEN et al., 1993). PcyA1 which is required for bilin synthesis, may play a critical role in assisting their folding, by modulating the activities of key enzymes of chlorophyll synthesis such as magnesium chelatase (MgCh) and light-dependent protochlorophyllide oxidoreductase (LPOR). Previous studies showed that CrPcyA1 physically interacts with LPOR (Zhang et al., 2018). Moreover, binding of PCB, the catalytic product of CrPcyA1, to GUN4 (genomes uncoupled 4) stimulates the enzymatic activity of MgCh (Zhang et al., 2021). The association of CrPcyA1 with the ATPase motor observed in our structure suggest that CrPcyA1 regulates the supply of chlorophylls to the newly imported unfolded proteins. In addition, bilins also participate in retrograde signaling in oxygenic photosynthetic organisms for the detoxification of reactive oxygen species generated in the light (Duanmu et al., 2013). Thus, the question arises whether the Ycf2-motor complex could mediate the transfer of the signal out of the chloroplast through the IEM which could then diffuse to the cytosol through the OEM known to be more permeable than the IEM. We also found that several proteins in the Orf2971-FtsHi complex are presumably involved in lipid/fatty acids (FAs) processing according to their structures. The mechanistic details of the lipid metabolism in plants have been extensively studied (Chan and Vogel, 2010; Cronan, 2014; Li-Beisson et al., 2015). The *de novo* synthesis of FAs in plants occurs in the chloroplast stroma. During synthesis, fatty acyl intermediates are covalently bound to ACPs, and the acyl-ACPs are then hydrolyzed into free FAs. While a portion of free FAs is utilized to produce chloroplast membrane lipids which contain unsaturated fatty acyl chains generated by FADs, a majority of free FAs is exported from the chloroplast into the cytosol and/or ER, presumably through FAXs located in the IEM of chloroplasts (Li-Beisson et al., 2017; Li et al., 2016) (Li-Beisson et al., 2013). We observed that in the Orf2971-FtsHi complex, the stroma-localized ACP2 is linked to the IEM-embedded proteins FAD6a-FaxL through Moc25, and together with Ctap7, an IMS located protein adopting a fatty acid binding protein folding, are positioned in close proximity to each other and at the periphery of the complex. These proteins might collaborate in lipid production and trafficking. The presence of these proteins in Orf2971-FtsHi motor may help to maintain the stability of lipids or membrane metabolism in a confined membrane region, thus avoiding membrane disturbances caused by frequent protein trafficking.

Finally it is interesting to note that both Orf2971 and Tic214 of the TOC-TIC complex which function as a scaffold and play a key role in the assembly of these two complexes are both encoded by the chloroplast genome in contrast to all the other nucleus-encoded subunits of these complexes. Whether the maintenance of the genes of Orf2971 and Tic214 in the chloroplast is linked to the assembly mechanism of these two complexes remains an intriguing question.

## Methods and materials

### Chlamydomonas strain construction and growth condition

The Orf2971-FLAG tagged strain was generated as previously described (Xing et al., 2022). In brief, the *orf2971’s* upstream and downstream regions fused with the FLAG sequence were amplified and cloned into the pUC-atpX-aadA vector. A31, a host strain graciously provided by Jean-David Rochaix, was utilized for the transformation process. The algal cells were cultured in TAP medium at 25 °C under continuous cool-white fluorescent light (60 µmol photons m^-2^ s^-1^) and harvested during the exponential growth phase.

### Isolation and purification of Orf2971-FtsHi motor complex

The cells of *Chlamydomonas reinhardtii* transgenic strain containing FLAG-tagged Orf2971 were collected through centrifugation. The pellet was suspended using Buffer A (25 mM Mops-KOH, pH 7.5, 200 mM NaCl, 10% Glycerol, and cocktail protease inhibitor solution (Roche) with 50-fold dilution), then lysed by high-pressure (500 Pa) for three times. The slurries were centrifuged at 15,000 g, 10 mins, 4 °C. The supernatant was further centrifuged at 200,000 g, 1 hour, 4 °C. The pellet containing plastid membranes was solubilized using Buffer B (25 mM Mops-KOH, pH 7.5, 100 mM NaCl, 10% Glycerol, 1% LMNG, and 0.12% CHS) for 1 hour. The solubilized membranes were centrifuged again and the dark green supernatant was incubated with FLAG-beads (Sigma-Aldrich) for 1 hour. The protein-loaded beads were washed with Buffer C (25mM Mops-KOH, pH 7.5, 100mM NaCl, 10% Glycerol, 0.0015% GDN) for five times, then eluted by Buffer D containing Buffer C and 200 µg/ml FLAG peptides for 1 hour. The elution containing the Orf2971-FtsHi complex was collected and concentrated to 1 mg/ml for cryo-sample preparation. The protein sample was analyzed by 4-20% gradient SDS-PAGE and Coomassie brilliant blue staining. The separated protein bands were respectively excised for further mass spectrometry analysis and data mining.

### Grid preparation and cryo-EM data acquisition

About 3 µl of the purified complex sample was applied to a freshly-glow discharged Au 300 mesh R1.2/1.3 grids coated with 2 nm carbon film (Quantifoil). The grids were blotted with force level of 1 and blot time of 4s, and flash plunged into the liquid ethane using Vitrobot (FEI) at 4 °C, 100% humidity, and then transferred into liquid nitrogen. Cryo-EM data were obtained on a 300 kV Titan Krios electron microscopy (Thermo Fisher Scientific), equipped with a K2 Summit camera (Gatan) and a quantum energy filter (GIF). Movies were recorded at a magnification of ×130,000, with a calibrated pixel size of 0.52 Å and a total dose of approximately 50 electrons per Å^2^ per 32 frames. The cryo-EM movies were obtained using SerialEM software (Mastronarde, 2005) with the defocus values ranged from -1.5 µm to -2.2 µm. A total of 10,634 movies were collected.

### Image processing

The movies stacks were motion corrected, dose-weighted, and binned to the pixel size of 1.04 Å, using MotionCorr2 (Zheng et al., 2017) in RELION 4.0 software (Scheres, 2012a, b). All calculating steps were performed using RELION 4.0 unless otherwise stated. The contrast transfer function was estimated by GCTF (Zhang, 2016) in RELION. The corrected images were imported to Gautomatch (https://github.com/JackZhang-Lab) for initial particle picking without any reference, which generated approximately 1,070,113 particles. The subsequent iterative-2D classification was applied to sieve the good particles for topaz training (Bepler et al., 2019). Next, the trained topaz model was used to obtain approximately 1,562,008 particles. Through several rounds of 2D classification and 3D classification, 127,613 particles with good quality were selected for auto3D refinement. After CTF refinement, Bayesian polishing and post-processing, the reconstruction map of Orf2971-FtsHi motor reached an overall resolution of 3.2 angstrom calculated using the gold standard Fourier shell correlation (FSC) of 0.143 criterion (Rosenthal and Henderson, 2003). We further performed several local refinements with specific masks to improve the map quality and resolution of local regions (Fig. S1C). All these local refined densities were combined into one composite map using volume maximum command in ChimeraX (Pettersen et al., 2021).

### Model building, refinement and validation

The motor complex model was built using the Cα backbone structure generated by DeepTracer (Pfab et al., 2021) or ModelAngelo (Jamali et al., 2022) as the initial reference. Combining the predicted structures by AlphaFold2 (Jumper et al., 2021; Mirdita et al., 2022) or ESMFold (Lin et al., 2023), all subunits had been manually assigned into the map using ChimeraX, according to the reliable secondary structure arrangements and sidechain density fitness. Several lipids and ligands were tentatively assigned and built based on their cryo-EM densities. The initial structural model of the motor complex was manually adjusted and refined against the composite sharpened map with COOT (Emsley and Cowtan, 2004). The resulted model was refined and validated with Phenix (Liebschner et al., 2019). The summaries for 3D reconstruction and model refinement were shown in Table S1. All figures of cryo-EM density and protein structure were produced by ChimeraX.

### ATPase activity assay

The ATPase activity of motor complex was determined by the ATP/NADH coupled enzyme assay that measures the production of NAD^+^ as a readout for ATP hydrolysis (Nørby, 1988). The coupled enzyme system contains two additional enzymes, namely the pyruvate kinase that uses ADP to convert phosphoenolpyruvate to pyruvate, and the lactate dehydrogenase that utilizes newly synthesized pyruvate to oxidize NADH to NAD^+^, which could be directly detected by the absorbance at 340 nm. In our experiments, all reactions were conducted at 30 °C in the buffer containing 50 mM Mops-KOH, pH7.4, 150 mM NaCl, 10 mM KCl, 10 mM MgCl_2_, 0.008% GDN, and in the presence of 1mM ATP and 0.025 µM motor complex, in 200µL reaction volume. In the control group setting, 0.025 µM motor complex was replaced by the equal volume of double distilled water. Absorbance was measured with a UV spectrophotometer (U-3900, Hitachi) using transparent quartz cell. The experiment was repeated three times and similar results were obtained. Data statics was performed in GraphPad Prism 8.

### Protein expression and purification

The genes encoding POTRA domains of Toc75 (Toc75-POTRA) and Tic22 were synthesized (GenScript, China) and constructed into the vector pET-21b with an N-terminal His-tag (Tic22) or an N-terminal GST-tag (Toc75-POTRA). The Toc75-POTRA vector also contains a tobacco etch virus (TEV) protease cleavage site downstream the GST-tag, which allows the removing of the GST tag from the mature protein. The recombinant plasmids were transformed into E. coli strain BL21 (DE3) (Transgen, China). The transformed cells were cultured in Lysogeny Broth medium containing 100 µg mL^−1^ Ampicillin and shaken at 37 °C until the O.D._600_ reaching around 1.5. The protein expression was induced by adding 1 mM IPTG (isopropylβ-D-1-thiogalactopyranoside), and the cells were further shaken at 16 °C for 18 h. The cells were harvested by centrifugation at 8000 g for 6 min. The cell pellets were resuspended in PBS buffer (20 mM, pH 7.4) and disrupted by ultrasonication. The cell lysate was centrifuged at 35,000 g for 30 min. To purify the Tic22 protein, the supernatant was loaded onto the Ni-sepharose fast flow resin (GE Healthcare), which was pre-equilibrated with PBS. After binding for about 1 h, the Ni-sepharose fast flow resin was washed with the PBS buffer containing 30 mM and 50 mM imidazole. The Tic22 protein was then eluted with PBS buffer containing 250 mM imidazole. To purify the Toc75-POTRA protein, the supernatant was loaded onto the glutathione-sepharose beads (GE Healthcare) and was washed with PBS buffer, and the Toc75-POTRA protein was eluted with PBS buffer containing 5mM glutathione. The eluted Toc75-POTRA protein was then digested with TEV protease and further purified using the glutathione-sepharose beads to remove GST tag. Both purified proteins were dialyzed in PBS buffer and then concentrated using an Ultracel centrifugal filter with a 10-kDa cutoff.

### SPR-based biosensor assay

The binding affinity between Tic22 and Toc75-POTRA was measured by SPR using a Biacore8K system and a CM5 chip (GE Healthcare). Toc75-POTRA was diluted with 10 mM NaAC, pH 4.5 to the concentration of 33 µg mL^−1^, and then immobilized on the activated CM5 chip using an amine coupling method. The running buffer contains 20 mM PBS (pH 7.4) with 0.05% Tween 20. The Tic22 proteins with gradient concentrations (1, 2, 4, 8 and 16 µM) were flowed over the chip surface with a flow rate of 30 µL min^-1^, bound for 60 s and dissociated for 180 s. The single-cycle binding kinetics were analyzed with Biacore8K evaluation software (GE Healthcare) by fitting to a 1:1 binding model.

### Rosetta ensemble docking and molecular dynamic simulation

Sequences of Tic22 and POTRA1-3 domain (POTRAs) of Toc75 of *Chlamydomonas reinhardtii* were retrieved from Phytozome v13 database by corresponding identities (Cre14.g625750 and Cre03.g175200). Since that Tic22 model from our cryo-EM structure was only partially built, we employed AlphFold2 to predict a complete model for Tic22 (Mirdita et al., 2022), which can be well aligned with our structural model. The model of POTRAs of Toc75 was isolated from TOC-TIC structure (PDB 7XZI) (Liu et al., 2023), and the gap regions absent in the structure were built by Rosetta3.13 package. Next, we used Gromacs version 2020 (Van Der Spoel et al., 2005), Amber14SB force field (Maier et al., 2015) and Tip3p water model to perform the conventional molecular dynamic (cMD) simulation to sample multiple diverse conformations of Tic22 and POTRAs, respectively. The detailed parameters for product cMD simulation are as following: The Lincs algorithm was used for hydrogen bond constraint (Hess et al., 1997); The system temperature was set at 300 K, and temperature coupling method was V-rescale with time constant 0.1 ps; The pressure coupling way was Parrinello-Raham with tau_p 2.0 ps (Parrinello and Rahman, 1981); The reference standard pressure was set to 1.0 bar; Electrostatic interactions were calculation using the Particle Mesh Ewald (PME) method with 1.2 nm cut-off (Essmann et al., 1995). Finally, we generated several 500 ns time long trajectories after system had been equilibrated. We extracted the unbound ensemble from equilibrated trajectories for Tic22 and POTRAs separately, by clustering with 1.5 angstrom RMSD cut-off. After this, we applied Rosetta flexible docking way (Chaudhury and Gray, 2008) to explore the best docking conformation of Tic22–POTRAs complex, using their unbound ensemble produced by cMD. The total docking number is set to 100,000.

Both the RMSD values of docking poses against the starting coordinates, and the respective docking scores were recorded in graphic plotting. Subsequently, we chose the Tic22-POTRAs complex conformation with the lowest docking score for the Gaussian-accelerated MD (GaMD) simulation (Miao et al., 2015).

GaMD is a state of art enhanced sampling MD simulation method (Miao et al., 2015; Pang et al., 2017) that was extensively used in protein conformation transition, protein folding and protein ligand recognizing field (Bhattarai and Miao, 2018; Miao et al., 2018; Miao and McCammon, 2016; Wang and Miao, 2019). In our experiment, the GaMD simulation was performed using AMBER20 (Case et al., 2005; Salomon-Ferrer et al., 2013) with FF19SB protein force field (Tian et al., 2019) and OPC water model (Izadi et al., 2014). The protonation state of Tic22-POTRAs complex model was detected by H++ server (Anandakrishnan et al., 2012) at pH 7.0 condition. The complex model was solvated by water model in a cubic box where protein have a least 10.0 Å distance to the periodicity boundary, and neutralized by 100 mM NaCl. The simulation system was energy minimized with steepest descent algorithm for 2500 steps, followed by conjugated gradient method for 2500 steps with heavy atom position restraints of 2.0 kcal^-1^Å^-2^. The energy-minimized system was heated from 0 K to 300 K in 500 ps. Particle Mesh Ewald (PME) method was employed for electrostatic interaction calculations. Non-bonded cut-off distance was set at 8.0 Å. The SHAKE algorithm(Ryckaert et al., 1977) was used to constrain hydrogen bond interactions. The langevin dynamics with collision frequency gamma_ln was set to 2.0 ps^-1^. The system pressure control used Berendsen method with 1.0 bar and the pressure relaxation time was 2.0 ps. Before the product GaMD, the system was equilibrated for 500 ps at NVT with position restraint of 1.0 kcal^-1^Å^-2^, followed by 2 ns at NPT without any restraint. Finally, the GaMD simulations were performed based on this equilibrated system, containing 10 ns conventional MD simulation for recording the potential statistics to acquire GaMD acceleration parameter values, a 12 ns equilibration after adding the dual-boost potential. Finally, three independent ∼1 µs GaMD production simulations with randomized initial atomic velocities. The average and the standard deviation (SD) of the system potential energies were recorded every 500 steps. The coordinates were saved every 5000 steps. The production trajectories were adjusted by CPPTRAJ (Roe and Cheatham III, 2013) to remove the rotation and translation of system, and were used for RMSD, RMSF, Cluster and hydrogen-bond static analysis.

## Data availability

The atomic coordinate of the Orf2971-FtsHi complex has been deposited in the Protein Data Bank with the accession code 8XKS. The composite cryo-EM map of the complex has been deposited in the Electron Microscopy Data Bank with accession codes EMDB-38424. All other data generated or analysed are available from the corresponding authors on reasonable request. Source data are provided with this paper.

## Acknowledgements

We thank B. Zhu, X. Huang, X. Li, L. Chen and other staff members at the Center for Biological Imaging (IBP, CAS) for their support in data collection; L. Niu and M. Zhang from IBP, CAS for mass spectrometry. We are grateful to Profs. Jean-David Rochaix and Zhenfeng Liu for in-depth discussion. We thank Torsten Juelich (University of Chinese Academy of Sciences) for linguistic assistance during the preparation of the article. The project is funded by the Strategic Priority Research Program of CAS (XDB37020101), the National Key R&D Program of China (2021YFA0910800), National Natural Science Foundation of China (31930064), Youth Innovation Promotion Association, Chinese Academy of Sciences (Y2022038), Regional Joint Key Projects of the National Foundation of China (U22A20445), and the Natural Science and Foundation of Shandong Province (ZR2023ZD30).

## Author contributions

W.Y. and M.L. conceived and coordinated the project; J.X., J.P. and H. C.constructed the Chlamydomonas strains; N.W. performed the complex purification, the cryo-EM map reconstruction, structure building and refinement with assistance from X.S. and L.S.; N.W. performed the MD simulation; L.S. purified the recombinant proteins and performed the SPR experiment; N.W., J.X., W.Y. and M.L. analyzed the data; N.W., J.X., W.Y. and M.L. wrote the manuscript; all authors discussed and commented on the results and the manuscript.

## Supplemental Information

**Supplemental Figure 1-15**

**Supplemental Table 1-2**

**Supplemental Movie 1-2**

